# Cooperation between Imp2p and Cdc15p imparts stiffness to the constricting contractile ring in fission yeast

**DOI:** 10.1101/2022.06.11.495753

**Authors:** Kimberly Bellingham-Johnstun, Blake Commer, Brié Levesque, Zoe L. Tyree, Caroline Laplante

## Abstract

The contractile ring must anchor to the plasma membrane and cell wall to transmit its tension to the plasma membrane. F-BAR domain containing proteins including Cdc15p and Imp2p in fission yeast are likely candidate anchoring proteins based on their mutant phenotypes. Cdc15p is a node component, links the actin bundle to the plasma membrane, recruits Bgs1p to the division plane, prevents contractile ring sliding, and contributes to the stiffness of the contractile ring. Less is known about Imp2p. We found that similarly to Cdc15p, Imp2p contributes to the stiffness of the contractile ring and assembles into protein clusters. Imp2p clusters contain ∼8 Imp2p dimers and depend on the actin network for their stability at the division plane. Importantly, Imp2p and Cdc15p reciprocally affect the amount of the other in the contractile ring indicating that the two proteins influence each other during cytokinesis and may explain their similar phenotypes.

## INTRODUCTION

The mechanism that anchors the contractile ring to the plasma membrane and the extracellular matrix is critical for generating the tension that drives cytokinesis. Although, the exact molecular composition of this mechanism remains unclear, the Fer/CIP4 homology Bin-Amphiphysin-Rvs (F-BAR) domain containing proteins Cdc15p and Imp2p are strong candidates to the anchoring mechanism in fission yeast. Cells mutant for *cdc15* or depleted of Cdc15p and cells with mutations in *imp2* exhibit a contractile ring sliding phenotype (Arasada and Pollard, 2014; Martin-Garcia *et al*., 2014; McDonald *et al*., 2015; Snider *et al*., 2017; Snider *et al*., 2020; Willet *et al*., 2021). In those cells, the ring slides away from where it assembled before the onset of constriction resulting in asymmetric cell division. In fission yeast, a contractile ring sliding off the division plane during cytokinesis is a hallmark phenotype of defective anchoring of the contractile ring. Presumably, poor anchoring of the contractile ring results in its displacement along the cell length when the ring is experiencing sufficient tension (Arasada and Pollard, 2014).

More is known about the role of Cdc15p as a putative protein anchor than Imp2p. There are two possible explanations for the sliding of the contractile ring in Cdc15p deficient cells. The first explanation is that Cdc15p itself may anchor the contractile ring to the plasma membrane by interacting directly with the plasma membrane and indirectly with the actin cytoskeleton via binding of Cdc12p and Myo2 molecules (composed of the myosin heavy chain Myo2p and essential and regulatory light chains Cdc4p and Rlc1p) (Fankhauser *et al*., 1995; Carnahan and Gould, 2003; Roberts-Galbraith *et al*., 2009; Laporte *et al*., 2011; Willet *et al*., 2015; McDonald *et al*., 2017; Willet *et al*., 2018). The second explanation is that Cdc15p supports the anchoring of the contractile ring by promoting the transport of Bgs1p from the Golgi to the division plane where Bgs1p produces the septum material and stabilizes the position of the contractile ring (Cortes *et al*., 2002; Arasada and Pollard, 2014). The role of Imp2p in supporting the connection of the contractile ring to the plasma membrane and cell wall remains unknown.

The structural and phenotypic similarities between *imp2* and *cdc15* have concealed their distinctive roles during cytokinesis. Cdc15p and Imp2p share similar structural features with an N-terminal plasma membrane binding F-BAR domain followed by an intrinsically disordered region (IDR) and a C-terminal protein-protein interaction Src Homology 3 (SH3) domain. Imp2p and Cdc15p have partially interchangeable F-BAR, IDR and SH3 domains (Roberts-Galbraith *et al*., 2010; McDonald *et al*., 2016; Lee *et al*., 2018; Mangione *et al*., 2019; Bhattacharjee *et al*., 2020; Magliozzi *et al*., 2020; Willet *et al*., 2021). Imp2p and Cdc15p even share SH3-domain binding partners including Fic1p and Sbg1p. Some differences between the two proteins include the distinct impact of their regulation by phosphorylation. A recent study found that while prolonged phosphorylation of the IDR of Cdc15p leads to aberrations in membrane binding, the phosphorylation of the IDR of Imp2p is constitutive and necessary for proper contractile ring anchoring (Willet *et al*., 2021). Finally, Cdc15p is involved in both cytokinesis and endocytosis, whereas Imp2p localizes only to the contractile ring suggesting that it only functions during cytokinesis (Demeter and Sazer, 1998; Arasada and Pollard, 2011).

Cdc15p is a component of cytokinesis nodes, large protein complexes distributed in a broad band around the cell center that coalesce to form the contractile ring (Wu *et al*., 2003; Wu *et al*., 2006; Moseley *et al*., 2009). Quantitative single molecule localization microscopy (SMLM) in live cells revealed the molecular organization of node proteins (Laplante *et al*., 2016b; Bellingham-Johnstun *et al*., 2021). Node proteins closest to the plasma membrane form the core of the node and include Cdc15p, the IQGAP homolog Rng2p and the tips of the Myo2 tails, whereas the motor heads of the Myo2 molecules fan into the cytoplasm, poised to bind a somewhat random network of actin filaments ∼60 nm away from the plasma membrane. The molecular organization of the node thus hints that these protein complexes may act as anchors that link the main bundle of actin filaments to the plasma membrane and the cell wall. Recently, we investigated the role of Cdc15p on the mechanical properties of the contractile ring. We used laser ablation to sever the contractile ring of cells depleted of Cdc15p and found that Cdc15p impacts the stiffness of the contractile ring suggesting that nodes bear the tension load of the contractile ring (Moshtohry *et al*., 2022).

Here, we investigate the molecular organization of Imp2p in the contractile ring and its impact on the mechanical properties of the contractile ring. Analysis of the recoil displacement of severed contractile ring tips after laser ablation in Δ*imp2* cells show that Imp2p contributes to the stiffness of the contractile ring similarly to Ccd15p. SMLM in live cells revealed that Imp2p clusters into complexes within the contractile ring similar to Cdc15p nodes. However, the Imp2p clusters are smaller than the Cdc15p nodes and each cluster contains ∼8 dimers of Imp2p whereas each node contains ∼12 dimers of Cdc15p. We measured that the contractile ring contains fewer Imp2p clusters than Cdc15p nodes suggesting that these two complexes may not associate into stable larger complexes. Yet, Imp2p and Cdc15p reciprocally affect the levels of each other in the contractile ring indicating that they somehow influence each other. Unlike Cdc15p nodes, Imp2p clusters depend on the actin network for their stable interaction with the plasma membrane at the division plane underlining an important difference between the two complexes.

## RESULTS

### 1 Imp2p impacts the stiffness of the constricting contractile ring

Cells lacking Imp2p exhibit multiple cytokinesis defects including contractile rings that slide away from the division plane, defective ring disassembly and defective septation leading to multiseptated cells (Figure 1A-D) (Demeter and Sazer, 1998; Martin-Garcia *et al*., 2014). Cells lacking Imp2p showed contractile rings that were offset from the cell center similarly to cells depleted of Cdc15p (Figure 1B). To confirm that off center rings assemble in the center of the cell and slide away rather than assemble off center, we measured ring sliding in a timelapse of wild-type and Δ*imp2* cells expressing mEGFP-Myo2p and stained with Fluorescent Brighther 28 (FB28) to highlight the cell wall. We used cells depleted of Cdc15p as a control. In Δ*imp2* cells, rings slide away from their position of assembly over time similar to the ring sliding phenotype observed in Cdc15p-depleted cells (Figure 1C and D).

**Figure 1.**
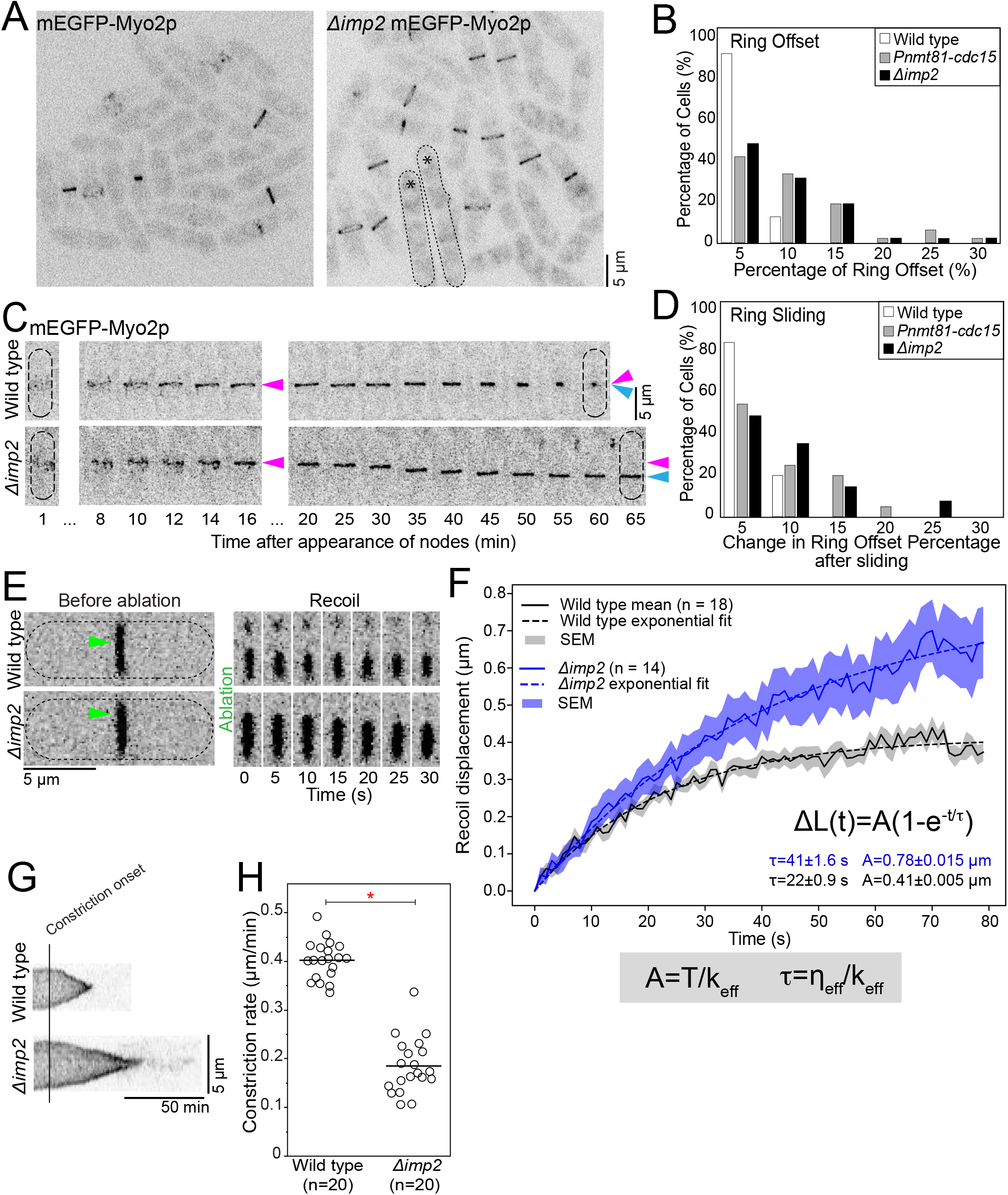
Imp2p impacts the stiffness of the constricting contractile ring. **A**. Confocal fluorescence micrographs of fields of wild-type (left) and Δ*imp2* cells (right). Asterisk, multiseptated cell. Dotted line, cell outline. **B**. Graph of the ring offset measured in still images. n = 50 cells for each genotype. **C**. Timelapse montages of wild-type and Δ*imp2* cells showing the position of the ring over time. Magenta arrowhead, position of the ring when the ring is assembled. Blue arrowhead, position of the ring at the end of cytokinesis (wild type) or maximum displacement (Δ*imp2*). n = 21 cells for each genotype. **D**. Graph of the change in ring offset measured after ring sliding in timelapse movies. **E**. Timelapse montages of wild-type (top) and Δ*imp2* (bottom) cells before and after laser ablation. Green arrowhead, site of ablation. **F**. Graph of the displacement of severed tips in wild-type and Δ*imp2* cells. Gray shaded box, equations representing our framework. **G**. Kymographs of constricting contractile ring aligned at constriction onset in a representative wild-type (top) and Δ*imp2* cell (bottom). **H**. Swarm plot showing the constriction rate of the contractile rings in wild-type and Δ*imp2* cells. Line, average constriction rate. Asterisk, p < 0.05 by Student’s t-test.

We recently described the contribution of Cdc15p to the stiffness of the constricting contractile ring (Moshtohry *et al*., 2022). Laser ablation of the contractile ring can reveal otherwise undetectable features about the structure including its mechanical properties (Silva *et al*., 2016; Moshtohry *et al*., 2022). We hypothesized that Imp2p may impact the mechanical properties of the ring in a distinct manner that would reveal specific functions in cytokinesis. We used laser ablation to sever constricting contractile rings in wild-type and Δ*imp2* cells expressing mEGFP-Myp2p to highlight the bundle of actin filaments in the ring (Supplemental Figure 1A and Methods) (Moshtohry *et al*., 2022). We focused on contractile rings that were 20-50% constricted to avoid confounding results with non-constricting contractile rings that may have distinct mechanical properties. We imaged each cell prior to laser ablation, severed the contractile ring, and imaged the response every second for up to ∼5 minutes.

After laser ablation of the constricting contractile ring in wild-type cells, the severed tips recoiled away from each other revealing a growing gap in the ring (Moshtohry *et al*., 2022). We interpreted this response as the local release of tension present in the constricting contractile ring prior to ablation (McDargh *et al*., 2021). Consistent with this interpretation, the rate of ring constriction slowed down, but constriction did not stop after laser ablation suggesting that severing the ring causes the local release of tension while tension is at least partially maintained away from the cut site (Moshtohry *et al*., 2022). Eventually, the severed tips stopped recoiling. We observed the same response in Δ*imp2* cells except that the severed tips recoiled farther from each other revealing a larger gap in the ring (Figure 1E).

To determine the impact of Imp2p on the mechanical properties of the contractile ring, we tracked the position of the severed tips during the recoil phase and calculated the recoil displacement as the distance of the severed tips from their initial position immediately after the cut over time. The recoil displacements of the severed tips in both wild-type and *Δimp2* cells followed an exponential profile as expected for a material with viscoelastic properties (Figure 1F) (Moshtohry *et al*., 2022) (Kumar *et al*., 2006; Colombelli *et al*., 2009; Silva *et al*., 2016; Roca-Cusachs *et al*., 2017). The mean recoil displacement of the severed tips Δ*L* fits to a single exponential, 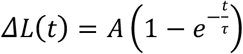. For severed rings in wild-type cells, we measured a total magnitude of the recoil after ablation, *A =* 0.41 ± 0.005 µm (mean ± standard error on the least squares fit, SE, n = 18 severed tips), and a timescale of viscoelastic recoil, *τ =* 22 ± 0.9 s (mean ± SE). For Δ*imp2* cells, both the recoil magnitude of the severed tips and timescale nearly doubled compared to wild type (*A*= 0.78 ± 0.015 µm, mean ± SE, n = 14) (*τ* = 41 ± 1.6 s, mean ± SE) (Figure 1F). The experimental displacement curve for *Δimp2* cells did not reach a plateau under our experimental conditions, likely because the severed tips recoil out of the imaging plane indicating that our reported values of *A* and *τ* may represent lower limits on these responses (Supplemental Figure 1A). The values of A and *τ* in Δ*imp2* cells are reminiscent of those measured in Cdc15p-depleted cells (*A* = 0.75 ± 0.013 µm and *τ* = 30 ± 1.2 s) (Supplemental Figure 1B) (Moshtohry *et al*., 2022).

We calculated the impact of Imp2p on the effective stiffness *k*_*eff*_ and effective viscous drag *η*_*eff*_of the contractile ring based on our framework for the contractile ring as a viscoelastic material under tension *T*_0_ (McDargh *et al*., 2021; Moshtohry *et al*., 2022). Both *A* and *τ* describe the displacement profile of the severed tips and are influenced by the mechanical properties of the contractile ring as *A* = *T*_0_/*k*_*eff*_ and *τ* = *η*_*eff*_ /*k*_*eff*_ (Figure 1F). Given the relationship between *A* and *k*_*eff*_ and the increased *A*, our data suggests that *k*_*eff*_ of the contractile ring decreases in Δ*imp2* cells. In addition, depleting Imp2p likely impacts *k*_*eff*_ more than *η*_*eff*_ as *A* and *τ* increased by a similar factor and both *A* and *τ* rely on *k*_*eff*_. We measured a reduction in the constriction rate in Δ*imp2* cells of ∼50% compared to wild type, suggesting that the total tension in the contractile ring is decreased in Δ*imp2* cells (Figure 1G and H). With the likelihood that *T*_0_ is decreased in the contractile rings of cells lacking Imp2p, the viscous drag may also be decreased compared to wild type. Nevertheless, deleting *imp2* impacts the stiffness more than the viscous drag.

### 2 Severed contractile rings in *Δimp2* cells heal by the recruitment of a strand of mEGFP-Myp2p

Following ablation, most severed contractile rings in both wild-type and Δ*imp2* cells are competent to heal the gap caused by the recoil (Figure 2A and B). In wild-type cells, the mEGFP-Myp2p signal filled the gap by 172 ± 67 min (n=9) after ablation. In Δ*imp2* cells, the healing took longer and the mEGFP-Myp2p signal filled the gap by 199 ± 56 min (n=8) after ablation. In wild-type cells, the severed tips stopped recoiling after 43 ± 21 s (n=18 severed tips) and in most cells the gap in the severed contractile ring healed over the following 82 ± 33 s (n=9 severed tips) (Moshtohry *et al*., 2022). Because most severed tips recoiled out of the imaging plane in Δ*imp2* cells, we could not determine when recoil ended or the duration of healing (Figure 2A and Supplemental Figure 1A). The slower healing in *Δimp2* cells may be due to their larger gap size.

**Figure 2.**
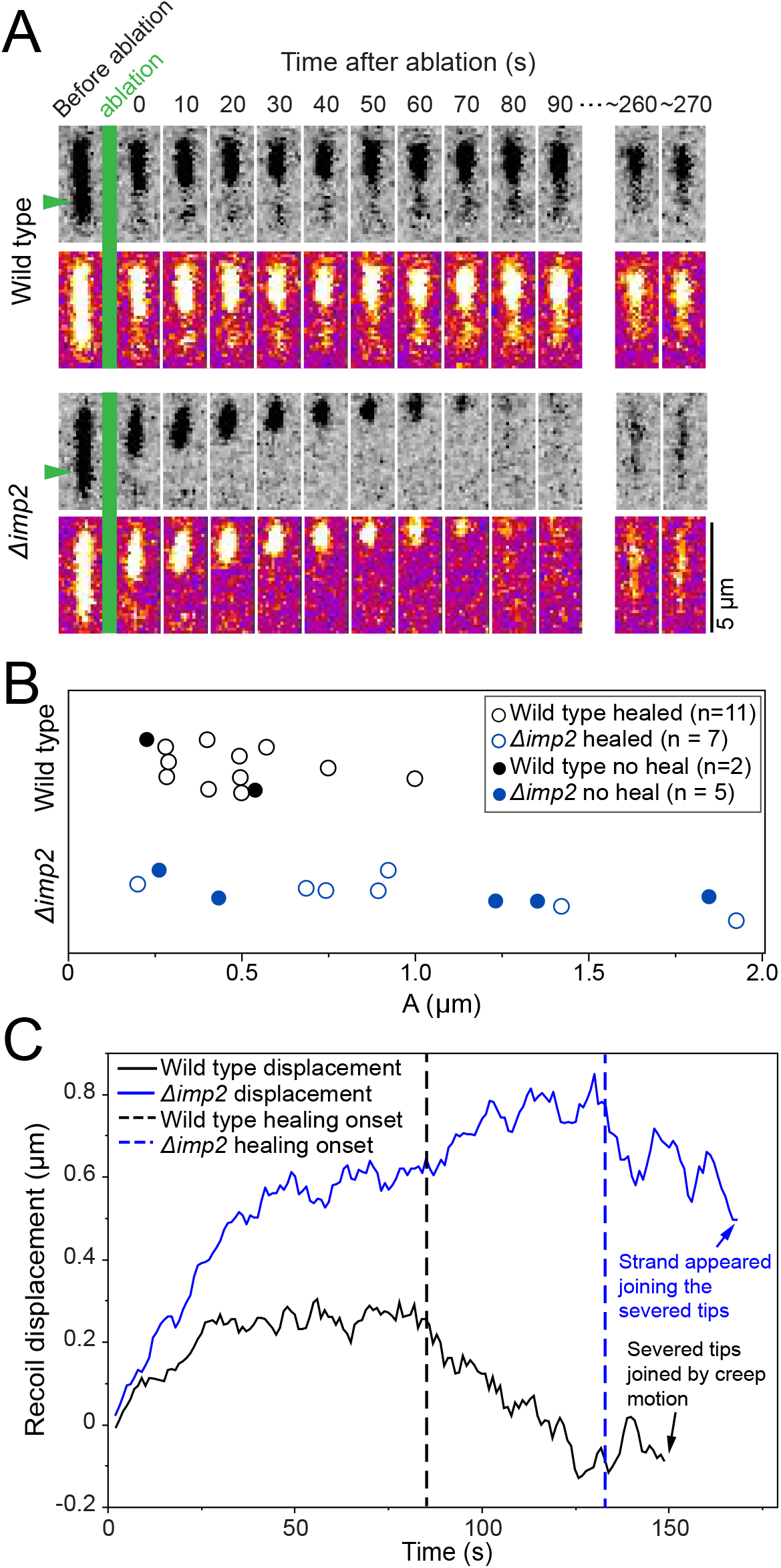
Severed contractile rings of *Δimp2* cells can heal despite a large gap size. **A**. Recoil and healing of typical severed wild-type (top) and Δ*imp2* (bottom) cells where the visible severed tip recoils out of the imaging plane. Top row, inverted gray LUT. Bottom row, Fire LUT. Green arrow, site of ablation. **B**. Graph of ability of a severed wild-type or Δ*imp2* contractile ring to heal after ablation versus maximum displacement. **C**. Graph of the displacement of a single severed tip of a wild-type or Δ*imp2* contractile ring showing recoil and healing. Dotted line, time at which healing stage begins. Tracking of the severed tips of wild-type ring show the progressive creeping motion until the tips. Tracking of the severed tips of Δ*imp2* ring shows initial progressive creeping motion of the tip until tracking ended when a strand appeared ∼170 s after ablation.

The healing process appeared different between wild-type and Δ*imp2* cells. Tracking the motions of the severed tips of wild-type rings show the progressive creeping motion until the tips connected (Figure 2A and C). In most severed rings of Δ*imp2* cells (5 out of 8 severed *Δimp2* rings healed) a strand labeled with mEGFP-Myp2p appeared in the plane of the ablated ring and filled the gap (Figure 2A and C). Tracking of the severed tips of Δ*imp2* rings showed initial progressive creeping motion of the tips until tracking ended when a strand appeared (Figure 2A and C and Supplemental Figure 2). The fluorescence intensity of the mEGFP-Myp2p signal in the strand then gradually increased over time (Figure 2A and Supplemental Figure 2). We observed this mechanism of healing severed rings with large and small gaps (Figure 2A and Supplemental Figure 2). The appearance of the strands suggests that a pre-assembled bundle of actin filaments labeled with mEGFP-Myp2p joined the discontinuous ring. The strands did not laterally sweep into the gap from within the imaging plane suggesting that they emerged from below the imaging plane. Although our current experimental setup limits acquisition to a single imaging plane preventing us from observing material across the thickness of the ring, dynamic bundles of actin filaments labeled with mEGFP-Myp2p cross the middle of the contractile ring during constriction and may be the source of the strands we observed during the healing of severed Δ*imp2* rings (Laplante *et al*., 2015).

### 3 Imp2p and Cdc15p affect the levels of each other in the contractile ring

To understand how Imp2p influences the mechanical properties of the contractile ring, we measured the local number of cytokinesis proteins in the contractile ring of Δ*imp2* cells and wild-type cells. We measured the number of polypeptides of type II myosins mEGFP-Myo2p and mEGFP-Myp2p, and of actin (labeled with *Pcof1-mEGFP-LifeAct*) in wild-type and *Δimp2* cells (Wu and Pollard, 2005; Wu *et al*., 2008; Malla *et al*., 2021). In brief, we acquired single timepoint z-stacks through the entire cell volume. We projected the images, corrected them for camera noise and uneven illumination and measured the local fluorescence intensity within the contractile ring for each mEGFP-tagged protein of interest. Finally, we use the local fluorescence intensity values to calculate the total number of polypeptides per contractile ring using a standard curve. We found no significant difference in the amount of either type II myosins or actin in the contractile ring of *Δimp2* cells compared to wild-type cells suggesting that Imp2p does not impact the tension of the contractile ring by altering the total amount of these proteins within the contractile ring (Figure 3A).

**Figure 3.**
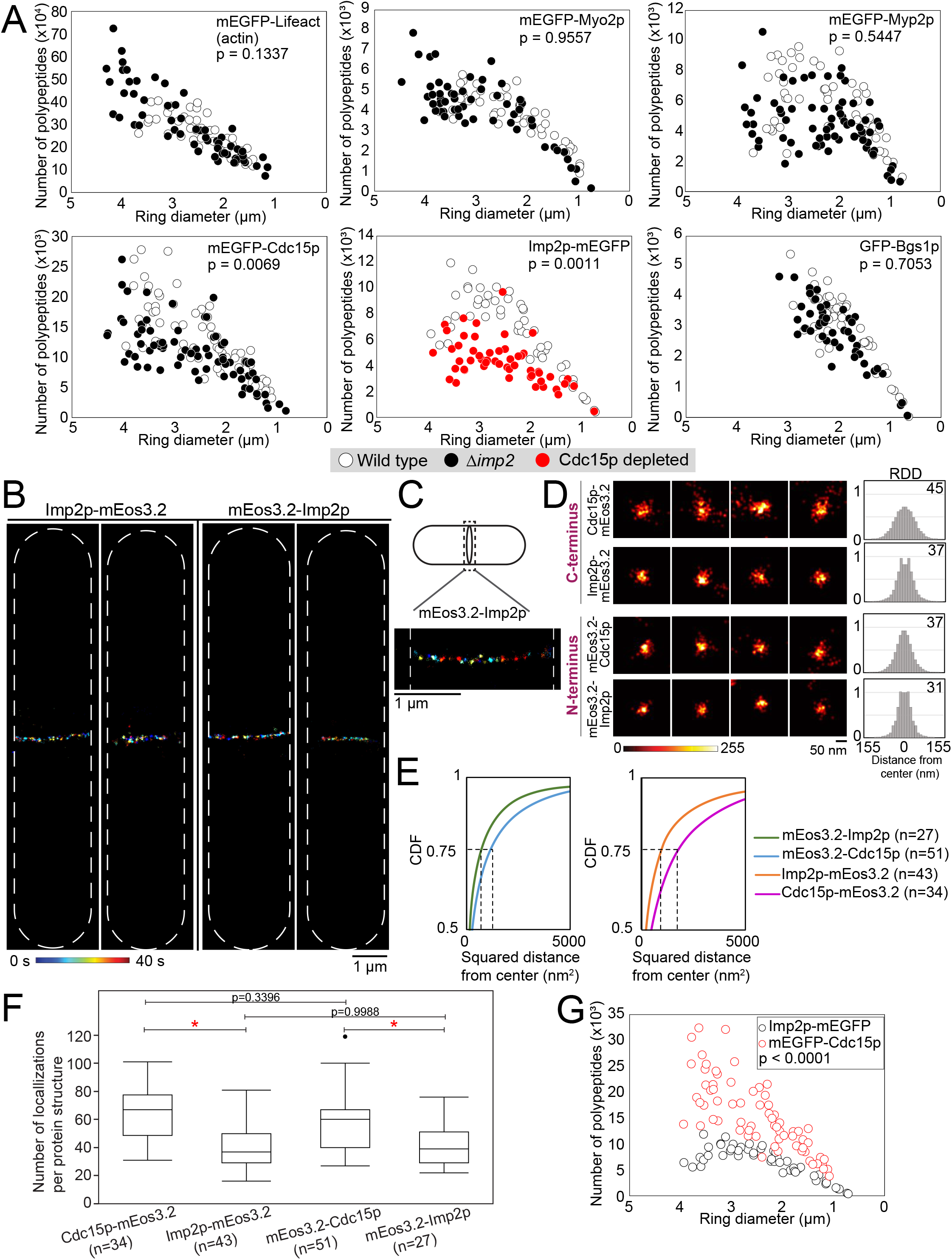
Imp2p can form clusters in the contractile ring. **A**. Plots of the distribution of the number of polypeptides per ring for cytokinesis proteins in wild-type, Δ*imp2*, and Cdc15p-depleted cells. Resulting actin monomer values from strains containing the mEGFP-Lifeact construct were scaled to reflect that the Lifeact construct labels 6% of polymerized actin (Malla *et al*., 2021). Differences between the distribution of polypeptides was determined by a Standard Least Squares model; p-values are stated on each plot. **B**. Examples of SMLM of Imp2p-mEos3.2 (left) and mEos3.2-Imp2p (right) in cells with contractile rings at different stage of constriction. **C**. Representative image of a cropped contractile ring in a cell expressing mEos3.2-Imp2p. **D**. Four representative cropped protein clusters from contractile rings of cells expressing either N- or C-terminal tagged Imp2p or Cdc15p (left). Protein clusters are color coded for density. Graphs of radial density distribution (RDD) and radii in nanometers (top right). **E**. Overlaid cumulative distribution function (CDF) plots of mEos3.2-Imp2p and mEos3.2-Cdc15p (left) and Imp2p-mEos3.2 and Cdc15p-mEos3.2 (right) in protein clusters of the contractile ring. Dotted line, squared distance away from center that contains 75% of localized emitters. Full CDF plots in Supplemental Figure 3B. **F**. Number of localized emitters per protein complex for N- or C-terminal tagged Imp2p or Cdc15p. Asterisk, p < 0.05 by Tukey’s HSD. **G**. Plots of the distribution of the number of polypeptides per ring for Imp2p-mEGFP and mEGFP-Cdc15p in wild-type cells. Differences between the distributions of polypeptides was determined by a Standard Least Squares model.

We determined whether the levels of Imp2p in the ring impact the levels of Cdc15p. We measured the levels of mEGFP-Cdc15p in Δ*imp2* and wild-type cells as described above. We measured a ∼25% decrease in the amount of Cdc15p in Δ*imp2* cells compared to wild type (Figure 3A). To determine whether Cdc15p affects the levels of Imp2p, we measured the local amount of Imp2p-mEGFP in the contractile rings of cells depleted of Cdc15p. We found that depleting Cdc15p caused a ∼50% decrease in the amount of Imp2p-mEGFP in contractile rings compared to wild type (Figure 3A). Therefore, our results suggest that in Δ*imp2* cells, there is ∼75% of the total amount of Cdc15p in the contractile ring. Whereas, in Cdc15p depleted cells, there is ∼3% of Cdc15p left and ∼50% of the total amount of Imp2p (Moshtohry *et al*., 2022).

Depleting Cdc15p results in a delayed accumulation of Bgs1p to the contractile ring and a decreased total amount of Bgs1p throughout constriction (Arasada and Pollard, 2014). Contractile ring sliding had been attributed to the decreased amount of Bgs1p in cells depleted of Cdc15p. As deleting *imp2* results in a ∼25% decrease of Cdc15p and mutations or deletion of *imp2* result in ring sliding, we sought to determine whether Imp2p impacts the amount of Bgs1p in the contractile ring by measuring the total amounts of GFP-Bgs1p in contractile rings of Δ*imp2* and wild-type cells (Willet *et al*., 2021). We found no significant difference in the total amount of GFP-Bgs1p in the rings of Δ*imp2* cells compared to wild type suggesting that although the lack of Imp2p causes a 25% decrease in Cdc15p, that decrease is insufficient to cause a noticeable decrease in Bgs1p-mEGFP. Our measurements suggest that contractile ring sliding in Δ*imp2* cells is not caused by a reduction in Bgs1p. Instead, the ring sliding phenotype may be caused by the combination of the lack of Imp2p and reduction of Cdc15p resulting in weak connections between the ring and the plasma membrane.

### 4 Complexes containing approximately 8 dimers of Imp2p assemble in the contractile ring

Our observation that Imp2p and Cdc15p reciprocally impact their levels in the ring suggest that they may assemble into a larger complex. Consistent with this hypothesis, SMLM of contractile rings in fixed cells showed that the N- and C-termini of Cdc15p and Imp2p localize in the same functional layers of the contractile ring (McDonald *et al*., 2017). Confocal imaging cannot distinguish the molecular organization of Imp2p and Cdc15p at the nanoscale in the contractile ring. Therefore, we used SMLM in live cells to determine the molecular organization of Imp2p in the constricting contractile ring. We tagged Imp2p at either the N- or the C-terminus with mEos3.2 and acquired SMLM data at 200 fps for 40 s (8,000 camera frames) (Laplante *et al*., 2016a; Laplante *et al*., 2016b; Bellingham-Johnstun *et al*., 2021). For comparison, we acquired both the N- and C-terminal mEos3.2 tagged Cdc15p. To enrich for cells in cytokinesis, we used *cdc25-22* mutation to arrest the cells at the G2-M phase of the cell cycle and released them into mitosis prior to imaging (see Methods). Cells expressing either the N- or C-terminal tagged Cdc15p showed the typical pattern of clustered emitters within the contractile ring (Laplante *et al*., 2016b). Similarly, cells expressing either of the N- or C-terminal mEos3.2 tagged Imp2p constructs showed clusters of emitters aligned into rings at the division plane suggesting that Imp2p assemble into protein complexes within the constricting contractile ring (Figure 3B and C).

To characterize the Imp2p protein clusters, we reconstructed the data using 1,000 camera frames (5 s), cropped the protein complexes and analyzed them to measure their radial density distribution (RDD) and dimensions (Figure 3D) (Laplante *et al*., 2016b; Bellingham-Johnstun *et al*., 2021). As controls, we measured the radii of mEos3.2-Cdc15p and Cdc15p-mEos3.2 nodes and obtained 37 nm and 45 nm respectively, comparable to our previous measurements (Laplante *et al*., 2016b; Bellingham-Johnstun *et al*., 2021). These measurements are consistent with the distribution of the N-terminal F-BAR domains grouping into smaller size clusters than the C-termini, located at the end of both the IDR and SH3 domains (Laplante *et al*., 2016b). We measured the radii of the N- and C-termini of Imp2p. Similarly, we found that the N-termini of Imp2p were smaller than their C-termini. Our measurements show a radius of 31 nm for mEos3.2-Imp2p and 37 nm for Imp2p-mEos3.2 (Figure 3D and E). Both N- and C-termini of Imp2p are smaller than those of Cdc15p (Figure 3D and E, and Supplemental Figure 3A and B).

We counted the number of localized emitters per cluster using the 5 s (1,000 camera frames) reconstructed data. We measured 58 ± 20 and 65 ± 19 localized emitters per mEos3.2-Cdc15p and Cdc15p-mEos3.2 nodes respectively. In contrast, we measured 42 ± 15 and 43 ± 17 localized emitters per mEos3.2-Imp2p and Imp2p-mEos3.2 cluster respectively (Figure 3F). Therefore, we measure ∼2/3 the total number of localized emitters for both Imp2p tagged constructs compared to the Cdc15p versions. Although the number of localized emitters per protein complex does not directly translate to number of proteins per protein complex due to the stochastic blinking behavior of mEos3.2, the number of emitters scales with the number of proteins per complex (Laplante *et al*., 2016b). In previous work, we estimated 10-12 dimers of Cdc15p per node, suggesting that there are ∼6-8 dimers of Imp2p per cluster (Laplante *et al*., 2016b).

We calculated the total number of Imp2p-mEGFP protein in the contractile ring by quantitative confocal microscopy as described above. In rings that were 20%-50% constricted, we calculated that the total amount of Imp2p-mEGFP is ∼1/2 of mEGFP-Cdc15p (Figure 3G). A ∼2/3 ratio of localized emitters for Imp2p to Cdc15p per node combined with a ∼1/2 ratio of total Im2p to Cdc15p per ring suggest that there are fewer Imp2p complexes than Cdc15p nodes per ring.

### 5 Imp2p requires actin for stable localization to the plasma membrane

The similarity in protein structure, localization within the contractile ring and impact on the mechanical properties in the contractile ring suggest that Cdc15p and Imp2p may be found in the same structure. Upon treating cells with a low concentration of Latrunculin A (LatA), nodes that were corralled in the contractile ring dispersed away from the division plane yet remained attached to the plasma membrane (Bellingham-Johnstun *et al*., 2021). To determine whether Cdc15p and Imp2p colocalize, we treated cells co-expressing Cdc15p-HALO*TMR and Imp2p-mEGFP with 10 µM LatA. We hypothesized that Imp2p clusters and Cdc15p nodes may disperse from the contractile ring either together or separately depending on their interactions. Within 10 minutes of LatA addition, Cdc15p-HALO*TMR nodes began to disperse away from the plane of cell division, as previously observed for nodes labeled with mEGFP-Myo2p (Figure 4A) (Bellingham-Johnstun *et al*., 2021). However, Imp2p-mEGFP did not exit the contractile ring as membrane-bound clusters. Instead, the Imp2p-mEGFP signal gradually disappeared from the contractile ring over a period of ∼50 min (Figure 4A and B). These observations suggested that Imp2p diffuses back into the cytoplasm as the actin of the contractile ring depolymerized. After ∼40 min of LatA treatment, we began to observe transient speckles of Imp2-mEGFP signal appearing at the plasma membrane throughout the cell surface and sometimes appearing adjacent to the nucleus (Figure 4A and B). These Imp2p-mEGFP accumulations were visible for only ∼ 1-3 timepoints (1-3 minutes) before disappearing. These observations suggest that the localization of Imp2p to the contractile ring and its interaction with the plasma membrane depend on the presence of actin.

**Figure 4.**
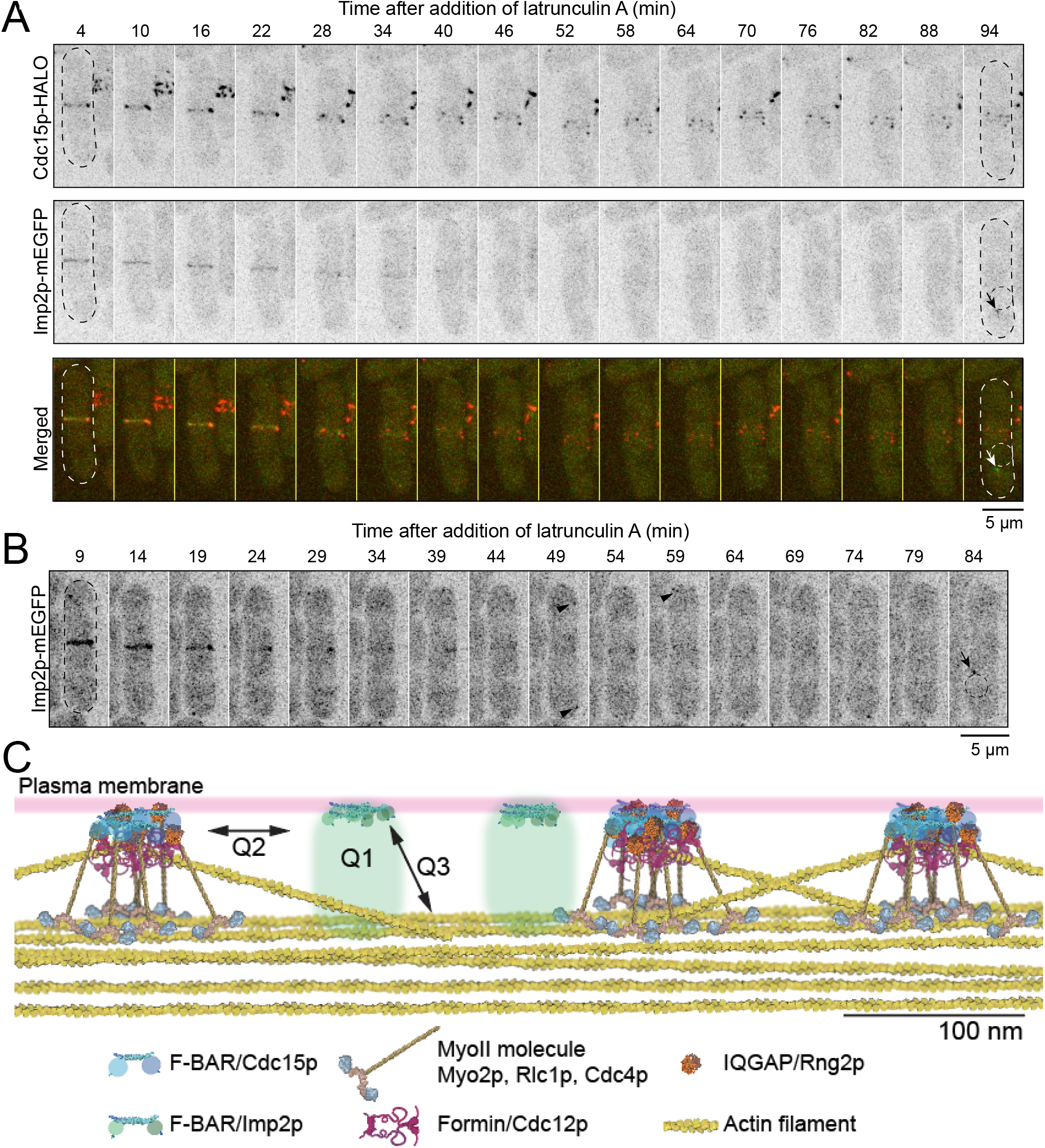
Imp2p localization at the plasma membrane is dependent on actin. **A**. Time series micrographs of cells co-expressing Cdc15p-HALO*TMR and Imp2p-mEGFP after the addition of 5 µM LatA. Arrow, perinuclear cluster of Imp2p. Dotted circular outline, nucleus. **B**. Time series micrographs of strains co- expressing Imp2p-mEGFP after the addition of 10 µM LatA. Arrow, perinuclear cluster of Imp2p. Arrowhead, speckles of Imp2p. Dotted circular outline, nucleus. **C**. Molecular models of the Imp2p complexes were built using the calculated node radii, RDD, stoichiometry of number of localizations, and available protein structures. Molecules are drawn to scale. Q1: What are the other protein components of the Imp2p complexes? Q2: How do the Imp2p complexes and Cdc15p nodes influence each other? Q3: How does the actin network influence the stability of Imp2p at the plasma membrane?

## DISCUSSION

The molecular organization of the contractile ring imparts its properties and governs its function, but these relationships remain unknown. The many similarities between *cdc15* and *imp2* have complicated our understanding of their specific functions during cytokinesis. Imp2p and Cdc15p share a similar protein domain architecture with an N-terminal F-BAR domain followed by an IDR and a C-terminal SH3 domain. Domain swapping experiments have shown these domains are at least partially interchangeable between Imp2p and Cdc15p (Roberts-Galbraith *et al*., 2010; Lee *et al*., 2018; Mangione *et al*., 2019; Bhattacharjee *et al*., 2020; Magliozzi *et al*., 2020). Their SH3 domains share at least some binding partners including Fic1p and Sbg1p that may bridge the functions of Cdc15p and Imp2p (Roberts-Galbraith *et al*., 2009; Bohnert and Gould, 2012; Ren *et al*., 2015; Sethi *et al*., 2016; Bhattacharjee *et al*., 2020).

We used SMLM in live fission yeast cells and laser ablation to reveal the organization of the putative anchor protein Imp2p and determine its impact on the mechanical properties of the contractile ring. We found that Imp2p can assemble into protein complexes and cooperate with Cdc15p nodes to impart stiffness to the constricting contractile ring. Severing constricting contractile rings in Δ*imp2* cells resulted in a recoil displacement profile strikingly similar to that of severed rings in cells depleted of Cdc15p with an approximate doubling of *A* and *τ* in both strains. Based on our framework, increasing both *A* and *τ* by a similar factor indicates a decrease in the stiffness of the contractile ring with little or no impact on the viscous drag. Therefore, our results suggest that deleting *imp2* or depleting Cdc15p each affect the stiffness of the constricting contractile ring by a similar magnitude. Our results also showed that deleting Imp2p causes a ∼25% decrease in Cdc15p while depleting Cdc15p down to ∼3% of its total cellular concentration results in a ∼50% decrease in Imp2p (Moshtohry *et al*., 2022). Therefore, the overall measured impact on the stiffness measured in *Δimp2* and Cdc15-depleted cells is caused by the combined reduction of both F-BAR proteins.

Our results suggest that Imp2p and Cdc15p primarily impact the effective stiffness of the constricting contractile ring (Moshtohry *et al*., 2022). Although they may also impact the viscous drag, that effect is minimal compared to the impact on the stiffness. This minimal impact on the viscous drag may be due to the large number of molecular interactions that contribute to the effective viscous drag, including molecular interactions between actin filaments and between the bundle of actin and the membrane. To explain the impact of Imp2p and Cdc15p on the effective stiffness of the contractile ring, we assume they function as a Hookean spring. Upon severing the ring by laser ablation, the tension is released locally, and the spring stretches to a maximum distance *A*. The stiffness of the spring determines that maximum distance. When subjected to the same amount of tension force, a stiffer spring stretches to a short maximum distance whereas a softer spring stretches to a greater maximum distance. Deleting *imp2* or depleting Cdc15p result in a softer spring that can stretch to a greater maximum distance. Assuming that Imp2p and Cdc15p are critical factors in the overall anchoring mechanism, a softer anchoring mechanism is presumably less effective at bearing the load of the constricting contractile ring. This may lead to inefficient transmission of contractile force from the actin bundle to the plasma membrane and cell wall resulting in a slower rate of constriction.

As observed in Δ*imp2* cells, severed rings in cells depleted of Cdc15p recoiled farther than severed rings in wild-type cells. However, severed rings in cells depleted of Cdc15p rarely healed while severed rings in Δ*imp2* cells were competent to heal similarly to wild types. In cells depleted of Cdc15p, the ability of a severed ring to heal diminished when the size of the gap created by the recoil was greater than ∼ 1 µm. This relationship between the ability of a gap to heal and the size of the gap is consistent with a healing mechanism based on the Search-Capture-Pull-Release mechanism limited to joining ring segments separated by less than ∼1 µm (Vavylonis *et al*., 2008; Moshtohry *et al*., 2022). In contrast, gaps of all sizes healed in Δ*imp2* cells indicating that the ability of a severed ring to heal did not correlate with the size of the gap in those cells (Figure 2B). Furthermore, although severed tips in Δ*imp2* cells can creep toward each other (creep healing method), severed rings healed by the appearance of a strand across the gap (strand capture healing method). The few severed rings that healed in cells depleted of Cdc15p healed by the creep healing method rather than the strand capture healing method. Our observations suggest that these two complementary methods can heal a discontinuous ring: the severed tip creep healing method employed by wild-type and Cdc15p-depleted cells and the strand capture healing method observed in Δ*imp2* cells. Severed rings in wild-type cells may also use the strand capture healing method but since the gaps created by the recoil are typically small the creep healing method may be sufficient to heal the severed ring without the need for an emerging strand. Alternatively, we may not be capable of distinguishing the arrival of an emerging strand once the gap size is diminished by the action of the creep healing method. The strands may originate from the center of the contractile ring below the imaging plane. Strands or actin bundles cross the center of the contractile ring during constriction (Laplante *et al*., 2015). Consistent with this hypothesis, we do not visualize strands sweeping into the gap of the contractile ring laterally. Cdc15p may be essential for capturing emerging strands as gaps in severed rings of Cdc15p depleted cells do not heal by the strand capture healing methods. One possible mechanism of strand capture healing may be that Cdc15p nodes can recover in the gap and the Myo2 in the nodes can capture an emerging strand. These two healing methods may represent complementary mechanisms that ensure the robustness of cytokinesis. Severed constricting contractile rings in *C. elegans* are also competent to heal indicating that the mechanisms that heal discontinuous contractile rings may be conserved across species (Silva *et al*., 2016).

Using SMLM in live fission yeast cells, we found that Imp2p can assemble into complexes within the contractile ring (Figure 4C). The Imp2p protein complexes contain ∼8 Imp2p dimers whereas nodes contain ∼12 Cdc15p dimers (Laplante *et al*., 2016b). Although cells with mutations in *imp2* or Δ*imp2* show the hallmarked ring sliding phenotype, how Imp2p complexes support anchoring remains unclear (Demeter and Sazer, 1998; Willet *et al*., 2021). To determine the function of the Imp2p complexes during cytokinesis, future work will need to answer three main questions (Figure 4C): 1) What are the other components of the Imp2p complex and how are they organized within the complex?, 2) What is the molecular relationship between Imp2p complexes and Cdc15p nodes that allow their reciprocal influence?, and 3) What is the nature of the link between the Imp2p complex and the actin network?

Understanding the molecular composition and organization of the Imp2p complex may reveal how they may anchor the ring during cytokinesis. Both Imp2p and Cdc15p localize within the same functional layer of the contractile ring consistent with their possible overlapping roles during cytokinesis (McDonald *et al*., 2017). Our more extensive knowledge of the molecular organization of Cdc15p nodes gave us clues about how nodes may act within the anchoring mechanism. Cdc15p is a node component that connects the contractile actin bundle, via interactions with Rng2p which binds Myo2p and Cdc12, to the plasma membrane and cell wall (Figure 4C) (Carnahan and Gould, 2003; Roberts-Galbraith *et al*., 2010; Laporte *et al*., 2011; Willet *et al*., 2015). The molecular organization of the nodes, the biochemical functions of node proteins and the ring sliding phenotype of cells expressing mutant *cdc15* or depleted of Cdc15p suggest that Cdc15p nodes act as ring anchors. Cdc15p likely influences the anchoring of the contractile ring by supporting the timely transport of Bgs1p to the division plane (Arasada and Pollard, 2014). Our measurements do not support a role for Imp2p in the recruitment of Bgs1p to the contractile ring.

Our quantitative measurements show that Imp2p and Cdc15p reciprocally affect the levels of each other. How can these two proteins influence the level of the other? One possibility is that Cdc15p nodes and Imp2p complexes physically interact with each other. For example, Imp2p complexes and Cdc15p nodes may be subunits of a large and stable megacomplex. Alternatively, the two complexes may interact transiently and dynamically. The nature of these molecular interactions is unknown but may be related to their shared binding partners Fic1p and Sbg1p. Fic1p and Sbg1p can interact with all three cytokinesis F-BAR domain containing proteins Cdc15p, Imp2p and Rga7p possibly relating these different components of the general anchoring mechanism (Roberts-Galbraith *et al*., 2009; Bohnert and Gould, 2012; Martin-Garcia *et al*., 2014; Ren *et al*., 2015; Sethi *et al*., 2016; Bhattacharjee *et al*., 2020). These shared physical interactions and other unknown ones may allow each protein complex to influence the other.

The stable association of Imp2p with the plasma membrane requires the presence of the actin network. In cells treated with LatA, the Imp2p-mEGFP signal originally in the contractile ring dispersed into the cytoplasm. Transient speckles of Imp2p-mEGFP then appeared at the plasma membrane in those cells suggesting that the F-BAR may be competent to bind the membrane in the absence of the actin network, but the actin network is required for increased stability of Imp2p at the plasma membrane. Currently, there are no known interacting partners of Imp2p that bind actin and Imp2p does not bind actin directly. Therefore, the nature of the relationship between Imp2p and the actin network remains unclear. Identifying other binding partners of Imp2p and other components of the Imp2p protein complex may reveal the nature of the dependency of Imp2p on actin.

## METHODS AND MATERIALS

### Strains, Growing Conditions, and Genetic and Cellular Methods

Table S1 lists the *S. pombe* strains used in this study. The strains were created using PCR-based gene targeting to integrate the constructs into the locus of choice and confirmed by PCR and fluorescence microscopy (Bahler *et al*., 1998). Either *pFA6a-(FPgene)-kanMX6* or *pFA6a-kanMX6-P(gene of interest)-mEos3*.*2* were used depending on whether C-terminal or N-terminal tagging of the gene was desired where (*FPgene*) denotes mEGFP, mEos3.2, mCherry or HALO. Primers with 80 bp of homologous sequence flanking the integration site (obtained at www.bahlerlab.info/resources/) and two repeats of GGA GGT to create a 4xGly linker were used to amplify the vector of choice. Except for the mEGFP-Lifeact construct, all tagged genes were under the control of their endogenous promoter. Cells were grown in an exponential phase for 36–48 hours before imaging. To deplete the expression of Cdc15p, cells with the *Pnmt81x-cdc15* or *Pnmt81x-mCherry-cdc15* construct were grown for 24 h in YE5S and switched to YE5S + 15 µM thiamine for 16-18 h prior to imaging.

Cells expressing Cdc15-HALO were stained with 250 nM of HaloTag® TMR Ligand, Cdc15p-HALO*TMR, (Promega, Catalog # G8251) for 1 h in a shaking incubator at 25°C. Cells were pelleted as described below. The supernatant was removed, and the pellet was rinsed three times with EMM5S medium. The cells were then washed for 30–60 min with EMM5S in a shaking incubator. Cells were then pelleted and mounted for imaging as described below.

To synchronize the population of cells, we used the temperature-sensitive *cdc25-22* mutation to arrest cells at the G2/M transition at the restrictive temperature of 34 °C for 4 h. We then released cells into mitosis at the permissive temperature of 22 °C as a synchronized population.

Treatment with LatA to depolymerize the actin cytoskeleton during imaging was performed with either 5 µM or 10 µM LatA. After addition of the drug, the cells were mounted as described below and imaged immediately.

Cells used to calculate ring offset and sliding were pelleted as described below and stained with Fluorescent Brightener 28 (FB28, Sigma-Aldrich) to highlight the cell wall. FB28 was stored at -20 °C in 5 mg/mL aliquots. Before use, aliquots were thawed and briefly spun down to remove debris. Immediately before imaging, 500 ug/mL FB28 was added to the sample and mixed gently before mounting as described below. A new aliquot was used for each imaging session.

### Spinning-Disk Confocal Microscopy and Data Analysis

Cells were grown in exponential phase at 25 °C in YE5S-rich liquid medium in 50-mL baffled flasks in a shaking incubator in the dark. Fluorescence images of live cells were acquired with a Nikon Eclipse Ti microscope equipped with a 100×/numerical aperture (NA) 1.49 HP Apo TIRF objective (Nikon), a CSU-X1 (Yokogawa) confocal spinning-disk system, 405/488/561/647 nm solid state lasers, and an electron-multiplying cooled charge-coupled device camera (EMCCD IXon 897, Andor Technology). The Nikon Element software was used for acquisition. Cells were concentrated 10- to 20-fold by centrifugation at 2,400 × *g* for 30 s and then resuspended in EMM5S. 5 μL of cells were mounted on a thin gelatin pad consisting of 10 μL 25% gelatin (Sigma-Aldrich; G-2500) in EMM5S, sealed under a #1.5 coverslip with VALAP (1:1:1 Vaseline:Lanolin:Parafin), and observed at 22 °C.

ImageJ (Schneider *et al*., 2012) and/or Nikon Element were used to create maximum or sum intensity projections of images, montages, and other image analyses. Except for time-lapse micrographs acquired after laser ablation, images in the figures are either maximum or sum intensity projections of z-sections spaced at 0.36 μm. Images were systematically contrasted to provide the best visualization, and images within the same figure panel were contrasted using the same settings. Confocal fluorescence micrographs in the figures are shown as inverted grayscale LUT. Ring constriction rate was measured using kymographs of maximum projection images (19 z-confocal planes taken for 6.48 μm) of time-lapse datasets taken at 1-min time intervals. The kymographs were thresholded, and the circumference was calculated automatically for each time point. These values were plotted in Microsoft Excel and the constriction rate calculated using a linear regression. Student’s t-tests were used to determine whether constriction rates differed significantly between planned comparisons of strains.

The percentage offset of the contractile was measured using maximum intensity projection images of timelapse of videos of cells expressing mEGFP-Myo2p and stained with FB28 to highlight the cell wall. The center of the cell was calculated as the cell length (L) divided by 2. The position of the contractile ring (X) was measured as the distance between the cell end and the contractile ring. For ring offset in still images, only rings that had septa stained with FB28 were used. The change in contractile ring offset was calculated using the following equation:

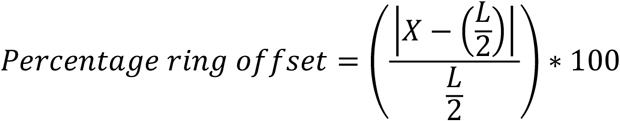

To calculate the percentage ring offset following sliding using timelapse micrographs, we measured the position of the contractile ring at the timepoint when it was fully assembled (X_a_) and the position of the contractile ring at the onset of constriction (X_c_). We calculated the difference between the ring offset at both timepoints using the following equation:

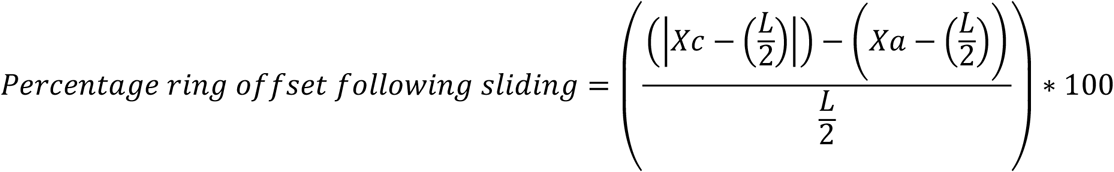

To count proteins in contractile rings, we created sum projection images of fields of cells from stacks of 21 optical images separated by 0.36 μm (Wu and Pollard, 2005; Wu *et al*., 2008). The images were corrected for the camera noise and uneven illumination, and then the fluorescence intensity of contractile rings was measured. These fluorescence intensity measurements were compared against a standard curve of proteins tagged endogenously with monomeric EGFP (mEGFP) to determine the number of molecules per contractile ring (Wu and Pollard, 2005; Wu *et al*., 2008). Resulting actin monomer values from strains containing the mEGFP-Lifeact construct were then scaled appropriately to reflect that this Lifeact construct marks 6% of polymerized actin (Malla *et al*., 2021). A Standard Least Squares model of the form Y = Ring Diameter + Genotype + Ring Diameter*Genotype, where Y is number of polypeptides, was used to determine whether the distribution of polypeptides in the constricting contractile ring differed between wild-type and either Δ*imp2* or *Pnmt81-cdc15* cells. Tests were performed with JMP Pro 16 (SAS Institute, Cary, NC).

### Super-resolution Data Acquisition and Display

Super-resolution imaging was performed with a Nikon STORM system operating in 2D mode, calibrated for single molecule acquisition in live cells. We used epi illumination to photoconvert and excite the fluorophores. We imaged single molecules with an sCMOS camera (ORCA-Flash4.0; Hamamatsu) operating at 200 frames per second using Nikon Element software. Powers for both 405 and 561 nm lasers were optimized for distinct minimally overlapping single molecule emissions (Bellingham-Johnstun *et al*., 2021). The average laser power density of the 561-nm laser used to excite the photoconverted mEos3.2 for imaging was ∼0.3 kW/cm^2^ illuminating a ∼5,500 µm^2^ area. To maintain a density of photoconverted mEos3.2 at an appropriate level for single-molecule localizations, the power of the 405-nm laser used for photoconversion was increased manually every 5 s during data acquisition. The total power of the 405-nm laser ranged from 0 to 32 µW.

Acquired data were processed to localize single molecules as previously described (Laplante *et al*., 2016a; Laplante *et al*., 2016b). Acquired frames were analyzed using a custom sCMOS-specific localization algorithm based on a maximum likelihood estimator (MLE) as described previously (Huang *et al*., 2013; Laplante *et al*., 2016a; Laplante *et al*., 2016b). A log-likelihood ratio was used as the rejection algorithm to filter out overlapping emitters, nonconverging fits, out-of-focus single molecules, and artifacts caused by rapid movements during one camera exposure time (Huang *et al*., 2011; Huang *et al*., 2013). The accepted estimates were reconstructed in a 2D histogram image of 5-nm pixels, where the integer value in each pixel represented the number of localizations estimates within that pixel. Images for visualization purposes were generated with each localization convolved with a 2D Gaussian kernel (σ = 7.5 nm). Images were reconstructed from all or a subset of acquired frames and color-coded for either the temporal information (JET LUT map) or for localization density (Heat LUT map). Our localization algorithm eliminated out-of-focus emissions, providing an effective depth of field of ∼400 nm (Laplante *et al*., 2016b).

### Node Identification and Measurements

Clusters of localized emitters associated with cytokinesis structures were manually selected from the reconstructed SMLM images. For comparison purposes, all nodes were cropped from images reconstructed from 1,000 frames (5 s) to minimize blurring due to crowding and movement and allow for the selection of individual nodes.

The edge of the cell was identified by increasing the brightness of SMLM images to enhance the cytoplasmic background. The edge of the cell was located where the cytoplasmic background drops off at the interface with the space outside the cells (Laplante *et al*., 2016b).

We analyzed ring nodes from contractile rings that had constricted by 20–50%. Ring nodes were difficult to segment if the contractile rings were constricted by over 50% due to the density and movement of the ring nodes. All nodes were cropped used a 309 × 309 nm box in MATLAB.

We used the spatial and temporal information provided by each localized mEos3.2 emitter to measure the dimensions and stoichiometry of proteins within ring nodes. By treating each localized emitter as an independent measurement and utilizing the large number of localizations, the radial density distribution approach becomes more robust than typical line profile measurements of fluorescence intensity.

Individual nodes were cropped, and the radial density distribution of their single molecule emitters were plotted as previously described (Laplante *et al*., 2016b). We reconstructed position estimates of the emitters in 2D histogram images from face views of isolated nodes obtained from the cropping step with pixel size of 2 nm to avoid pixelation errors in the subsequent measurement analysis. These images were then fit with a rotationally symmetric 2D Gaussian model with amplitude and sigma and center position in *x* and *y* as the fitting parameters. The radial symmetry centers of each node (Parthasarathy, 2012) were determined and used as the initial guesses for the *x, y* center in the fitting. MLE-based regression was performed assuming a Poisson noise model of the 2D histogram image using the Nelder–Mead simplex algorithm implemented in the MATLAB “fminsearch” function. Fitting estimates were filtered by their likelihood values thereafter. Fitting results that did not converge properly and resulted in center positions outside the image boundary or extremely large or small sigma values were also discarded from the results. This step eliminates instances where two objects were included in the same 309 × 309 nm selection box and include only selections containing a single cluster. For each node accepted through the filtering process, the distances of individual localizations from the estimated node center were calculated. All distances measured from a specific node type and view were plotted into histograms and then subsequently normalized by their radius to give a radial density distribution. The number of localizations per identified node was recorded as well.

We used a two-sample KS test to compare the distribution of localizations in each pair of node proteins. The null hypothesis is that the samples are drawn from the same distribution of localized emitters. The CDFs of the squared radial distance were calculated for each of the samples (Supplemental Figure 3B), and the maximum difference between pairs of CDFs was calculated and compared to the KS test critical value at a significance level of p < 0.005. If the maximum difference between the CDFs was greater than the critical value, the null hypothesis was rejected. The results of the KS test comparisons and sample size of the super-resolution datasets can be found in Supplemental Figure 3A.

The radius of each node marker was calculated using the CDF plots and was defined as the distance from the center of the node that contained 75% of the localized emitters (Figure 3E and Supplemental Figure 3B).

### Quantification of Localized Emitters

For each marker and genetic background, we measured the total number of localized emitters per protein complex. The number of localized emitters per complex is influenced by many factors, including the total number of frames used for the reconstruction, the photophysics of the fluorescent proteins, the number of tagged proteins per node, and the autofluorescent background.

We used an ANOVA with Tukey’s HSD test to compare the number of localizations per complex between genotypes at a significance level p < 0.05.

### Laser ablation and displacement analysis

We severed constricting contractile rings in wild-type and *Δimp2* cells expressing mEGFP-Myp2p. We focused on constricting contractile rings (∼75% of their initial size corresponding to 8-9 µm in circumference) to avoid potentially confounding results from contractile rings that have not yet started to constrict as they may have distinct mechanical properties. While we were able to visualize both severed tips in a few severed contractile rings, only one tip remained in our observation plane in most cases (Supplemental Figure 1A). We ablated 18 contractile rings in wild-type cells and 14 contractile rings in Δ*imp2* cells. This resulted in 18 severed tips that could be analyzed for the recoil phase of wild-type cells and 14 severed tips in Δ*imp2* cells.

Laser ablation was performed on a Nikon Ti-E microscope equipped with on an Andor Dragonfly spinning disk confocal fluorescence microscope equipped with 100x NA 1.45 objective (Nikon) and a built-in 1.5x magnifier; 488 nm diode laser with Borealis attachment (Andor); emission filter Chroma ET525/50m; and an EMCCD camera (iXon3, Andor Technology). Fusion software (Andor) was used to control data acquisition. Targeted laser ablation was performed using a MicroPoint (Andor) system with galvo-controlled steering to deliver 30 3 ns pulses of 551 nm light at 16 Hz (Andor) mounted on the Dragonfly microscope described above, as previously described (Moshtohry *et al*., 2022). Fusion software (Andor) was used to control acquisition while IQ software (Andor) was used simultaneously to control laser ablation. At this pulse rate, the ablation process lasts ∼2 s. Chroma ET610LP mounted in the dichroic position of a Nikon filter turret was used to deliver the ablation laser to the sample. Since this filter also reflects the mEGFP emission, the camera frames collected during the ablation process are blank. The behavior of severed contractile rings was imaged immediately following laser ablation by acquiring a single confocal plane in the 488-nm channel every second for up to 5 minutes. We acquired time-lapse images of a single optical plane of the surface of the contractile ring to maximize temporal resolution and reduce photodamage. The mechanism of ablative photodecomposition remains unclear but may be caused by either the propagation of a pressure wave and/or cavitation bubble dynamics (Venugopalan *et al*., 2002; Rau *et al*., 2006). The size of the damage is approximately the size of the diffraction spot of the lens <0.4 µm in XY plane and <0.8 in the Z axis (Khodjakov *et al*., 1997).

The displacement of the tips of severed contractile rings was tracked manually every second after laser ablation using the “multi-point” tool in FIJI on single z-plane time-lapse micrographs. The severed tips were tracked until they recoiled out of the imaging plane or until the contractile ring was healed. The coordinates of the tracked severed tips were exported as a CSV file. The displacement of the severed tips starting from the first frame after severing, t = 0 s, was calculated from the coordinates of the tracked severed tips using custom Python codes. The displacement traces from each cell were aligned to t = 0 s and the mean displacement curve was calculated for each genotype. The displacement traces show the combined recoil phase of all severed tips exclusively. The recoil displacement of each of the severed tips contributes to the traces until the recoil halts.

The mean displacement curve for each genotype was fit to a single exponential 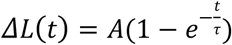 using a least squares fit, and the standard error on the least squares fit is reported. Contractile rings were considered fully ablated if there was no remaining fluorescence joining the ablated tips of the ring and the ablated ends showed evidence of recoil away from the site of ablation. Fully ablated rings also healed by the ablated tips slowly growing back towards each other, while the healing of partially ablated rings occurred as a uniform fluorescence recovery across the gap. This also resulted in the healing duration of partially ablated rings being faster than fully ablated rings. Ablated rings with an individual R_2_ < 0.3 were not included in Figure 2B due to poorness of fit.

## CONTRIBUTIONS

CL: Conceptualization. Methodology. Writing and editing. Project administration. Funding.

KBJ and BL: Data acquisition, analysis, and validation. Writing and editing.

BC: Data acquisition and analysis. Writing and editing.

ZT: Data analysis, validation, and editing.

## ACKNOWLEDGEMENTS

We thank Pilar Perez Gonzalez and Juan Carlos Ribas for sharing materials. We thank Parsa Zareiesfandabadi, Mary Williard Elting and Marcus Begley for technical support with laser ablation. We thank the CMIF at NCSU (supported by the State of North Carolina and NSF) for the ablation microscope. BC was supported by NIH T32 5T32-AI0502080-17. This work was funded by NIH R01GM134254.

## REFERENCES

Arasada, R., and Pollard, T.D. (2011). Distinct roles for F-BAR proteins Cdc15p and Bzz1p in actin polymerization at sites of endocytosis in fission yeast. Curr Biol 21, 1450–1459.

Arasada, R., and Pollard, T.D. (2014). Contractile ring stability in S. pombe depends on F-BAR protein Cdc15p and Bgs1p transport from the Golgi complex. Cell Rep 8, 1533–1544.

Bahler, J., Wu, J.Q., Longtine, M.S., Shah, N.G., McKenzie, A., 3rd, Steever, A.B., Wach, A., Philippsen, P., and Pringle, J.R. (1998). Heterologous modules for efficient and versatile PCR-based gene targeting in Schizosaccharomyces pombe. Yeast 14, 943–951.

Bellingham-Johnstun, K., Anders, E.C., Ravi, J., Bruinsma, C., and Laplante, C. (2021). Molecular organization of cytokinesis node predicts the constriction rate of the contractile ring. J Cell Biol 220.

Bhattacharjee, R., Mangione, M.C., Wos, M., Chen, J.S., Snider, C.E., Roberts-Galbraith, R.H., McDonald, N.A., Presti, L.L., Martin, S.G., and Gould, K.L. (2020). DYRK kinase Pom1 drives F-BAR protein Cdc15 from the membrane to promote medial division. Mol Biol Cell 31, 917–929.

Bohnert, K.A., and Gould, K.L. (2012). Cytokinesis-based constraints on polarized cell growth in fission yeast. PLoS Genet 8, e1003004.

Carnahan, R.H., and Gould, K.L. (2003). The PCH family protein, Cdc15p, recruits two F-actin nucleation pathways to coordinate cytokinetic actin ring formation in Schizosaccharomyces pombe. J Cell Biol 162, 851–862.

Colombelli, J., Besser, A., Kress, H., Reynaud, E.G., Girard, P., Caussinus, E., Haselmann, U., Small, J.V., Schwarz, U.S., and Stelzer, E.H. (2009). Mechanosensing in actin stress fibers revealed by a close correlation between force and protein localization. J Cell Sci 122, 1665–1679.

Cortes, J.C., Ishiguro, J., Duran, A., and Ribas, J.C. (2002). Localization of the (1,3)beta-D-glucan synthase catalytic subunit homologue Bgs1p/Cps1p from fission yeast suggests that it is involved in septation, polarized growth, mating, spore wall formation and spore germination. J Cell Sci 115, 4081–4096.

Demeter, J., and Sazer, S. (1998). imp2, a new component of the actin ring in the fission yeast Schizosaccharomyces pombe. J Cell Biol 143, 415–427.

Fankhauser, C., Reymond, A., Cerutti, L., Utzig, S., Hofmann, K., and Simanis, V. (1995). The S. pombe cdc15 gene is a key element in the reorganization of F-actin at mitosis. Cell 82, 435–444.

Huang, F., Hartwich, T.M., Rivera-Molina, F.E., Lin, Y., Duim, W.C., Long, J.J., Uchil, P.D., Myers, J.R., Baird, M.A., Mothes, W., Davidson, M.W., Toomre, D., and Bewersdorf, J. (2013). Video-rate nanoscopy using sCMOS camera-specific single-molecule localization algorithms. Nat Methods 10, 653–658.

Huang, F., Schwartz, S.L., Byars, J.M., and Lidke, K.A. (2011). Simultaneous multiple-emitter fitting for single molecule super-resolution imaging. Biomed Opt Express 2, 1377–1393.

Khodjakov, A., Cole, R.W., and Rieder, C.L. (1997). A synergy of technologies: combining laser microsurgery with green fluorescent protein tagging. Cell Motil Cytoskeleton 38, 311–317.

Kumar, S., Maxwell, I.Z., Heisterkamp, A., Polte, T.R., Lele, T.P., Salanga, M., Mazur, E., and Ingber, D.E. (2006). Viscoelastic retraction of single living stress fibers and its impact on cell shape, cytoskeletal organization, and extracellular matrix mechanics. Biophys J 90, 3762–3773.

Laplante, C., Berro, J., Karatekin, E., Hernandez-Leyva, A., Lee, R., and Pollard, T.D. (2015). Three myosins contribute uniquely to the assembly and constriction of the fission yeast cytokinetic contractile ring. Curr Biol 25, 1955–1965.

Laplante, C., Huang, F., Bewersdorf, J., and Pollard, T.D. (2016a). High-Speed Super-Resolution Imaging of Live Fission Yeast Cells. Methods Mol Biol 1369, 45–57.

Laplante, C., Huang, F., Tebbs, I.R., Bewersdorf, J., and Pollard, T.D. (2016b). Molecular organization of cytokinesis nodes and contractile rings by super-resolution fluorescence microscopy of live fission yeast. Proc Natl Acad Sci U S A 113, E5876–E5885.

Laporte, D., Coffman, V.C., Lee, I.J., and Wu, J.Q. (2011). Assembly and architecture of precursor nodes during fission yeast cytokinesis. J Cell Biol 192, 1005–1021.

Lee, M.E., Rusin, S.F., Jenkins, N., Kettenbach, A.N., and Moseley, J.B. (2018). Mechanisms Connecting the Conserved Protein Kinases Ssp1, Kin1, and Pom1 in Fission Yeast Cell Polarity and Division. Curr Biol 28, 84–92 e84.

Magliozzi, J.O., Sears, J., Cressey, L., Brady, M., Opalko, H.E., Kettenbach, A.N., and Moseley, J.B. (2020). Fission yeast Pak1 phosphorylates anillin-like Mid1 for spatial control of cytokinesis. J Cell Biol 219.

Malla, M., Pollard, T.D., and Chen, Q. (2021). Counting actin in contractile rings reveals novel contributions of cofilin and type II myosins to fission yeast cytokinesis. Mol Biol Cell, mbcE21080376.

Mangione, M.C., Snider, C.E., and Gould, K.L. (2019). The intrinsically disordered region of the cytokinetic F-BAR protein Cdc15 performs a unique essential function in maintenance of cytokinetic ring integrity. Mol Biol Cell 30, 2790–2801.

Martin-Garcia, R., Coll, P.M., and Perez, P. (2014). F-BAR domain protein Rga7 collaborates with Cdc15 and Imp2 to ensure proper cytokinesis in fission yeast. J Cell Sci 127, 4146–4158.

McDargh, Z., Wang, S., Chin, H.F., Thiyagarajan, S., Karatekin, E., Pollard, T.D., and O’Shaughnessy, B. (2021). Myosins generate contractile force and maintain organization in the cytokinetic contractile ring. bioRxiv, 2021.2005.2002.442363.

McDonald, N.A., Lind, A.L., Smith, S.E., Li, R., and Gould, K.L. (2017). Nanoscale architecture of the Schizosaccharomyces pombe contractile ring. Elife 6.

McDonald, N.A., Takizawa, Y., Feoktistova, A., Xu, P., Ohi, M.D., Vander Kooi, C.W., and Gould, K.L. (2016). The Tubulation Activity of a Fission Yeast F-BAR Protein Is Dispensable for Its Function in Cytokinesis. Cell Rep 14, 534–546.

McDonald, N.A., Vander Kooi, C.W., Ohi, M.D., and Gould, K.L. (2015). Oligomerization but Not Membrane Bending Underlies the Function of Certain F-BAR Proteins in Cell Motility and Cytokinesis. Dev Cell 35, 725–736.

Moseley, J.B., Mayeux, A., Paoletti, A., and Nurse, P. (2009). A spatial gradient coordinates cell size and mitotic entry in fission yeast. Nature 459, 857–860.

Moshtohry, M., Bellingham-Johnstun, K., Elting, M.W., and Laplante, C. (2022). Laser ablation reveals the impact of Cdc15p on the stiffness of the contractile ring. Mol Biol Cell 33, br9.

Parthasarathy, R. (2012). Rapid, accurate particle tracking by calculation of radial symmetry centers. Nat Methods 9, 724–726.

Rau, K.R., Quinto-Su, P.A., Hellman, A.N., and Venugopalan, V. (2006). Pulsed laser microbeam-induced cell lysis: time-resolved imaging and analysis of hydrodynamic effects. Biophys J 91, 317–329.

Ren, L., Willet, A.H., Roberts-Galbraith, R.H., McDonald, N.A., Feoktistova, A., Chen, J.S., Huang, H., Guillen, R., Boone, C., Sidhu, S.S., Beckley, J.R., and Gould, K.L. (2015). The Cdc15 and Imp2 SH3 domains cooperatively scaffold a network of proteins that redundantly ensure efficient cell division in fission yeast. Mol Biol Cell 26, 256–269.

Roberts-Galbraith, R.H., Chen, J.S., Wang, J., and Gould, K.L. (2009). The SH3 domains of two PCH family members cooperate in assembly of the Schizosaccharomyces pombe contractile ring. J Cell Biol 184, 113–127.

Roberts-Galbraith, R.H., Ohi, M.D., Ballif, B.A., Chen, J.S., McLeod, I., McDonald, W.H., Gygi, S.P., Yates, J.R., 3rd, and Gould, K.L. (2010). Dephosphorylation of F-BAR protein Cdc15 modulates its conformation and stimulates its scaffolding activity at the cell division site. Mol Cell 39, 86–99.

Roca-Cusachs, P., Conte, V., and Trepat, X. (2017). Quantifying forces in cell biology. Nat Cell Biol 19, 742–751.

Schneider, C.A., Rasband, W.S., and Eliceiri, K.W. (2012). NIH Image to ImageJ: 25 years of image analysis. Nat Methods 9, 671–675.

Sethi, K., Palani, S., Cortes, J.C., Sato, M., Sevugan, M., Ramos, M., Vijaykumar, S., Osumi, M., Naqvi, N.I., Ribas, J.C., and Balasubramanian, M. (2016). A New Membrane Protein Sbg1 Links the Contractile Ring Apparatus and Septum Synthesis Machinery in Fission Yeast. PLoS Genet 12, e1006383.

Silva, A.M., Osorio, D.S., Pereira, A.J., Maiato, H., Pinto, I.M., Rubinstein, B., Gassmann, R., Telley, I.A., and Carvalho, A.X. (2016). Robust gap repair in the contractile ring ensures timely completion of cytokinesis. J Cell Biol 215, 789–799.

Snider, C.E., Chandra, M., McDonald, N.A., Willet, A.H., Collier, S.E., Ohi, M.D., Jackson, L.P., and Gould, K.L. (2020). Opposite Surfaces of the Cdc15 F-BAR Domain Create a Membrane Platform That Coordinates Cytoskeletal and Signaling Components for Cytokinesis. Cell Rep 33, 108526.

Snider, C.E., Willet, A.H., Chen, J.S., Arpag, G., Zanic, M., and Gould, K.L. (2017). Phosphoinositide-mediated ring anchoring resists perpendicular forces to promote medial cytokinesis. J Cell Biol 216, 3041–3050.

Vavylonis, D., Wu, J.Q., Hao, S., O’Shaughnessy, B., and Pollard, T.D. (2008). Assembly mechanism of the contractile ring for cytokinesis by fission yeast. Science 319, 97–100.

Venugopalan, V., Guerra, A., 3rd, Nahen, K., and Vogel, A. (2002). Role of laser-induced plasma formation in pulsed cellular microsurgery and micromanipulation. Phys Rev Lett 88, 078103.

Willet, A.H., Bohnert, K.A., and Gould, K.L. (2018). Cdk1-dependent phosphoinhibition of a formin-F-BAR interaction opposes cytokinetic contractile ring formation. Mol Biol Cell 29, 713–721.

Willet, A.H., Igarashi, M.G., Chen, J.S., Bhattacharjee, R., Ren, L., Cullati, S.N., Elmore, Z.C., Roberts-Galbraith, R.H., Johnson, A.E., Beckley, J.R., and Gould, K.L. (2021). Phosphorylation in the intrinsically disordered region of F-BAR protein Imp2 regulates its contractile ring recruitment. J Cell Sci 134.

Willet, A.H., McDonald, N.A., Bohnert, K.A., Baird, M.A., Allen, J.R., Davidson, M.W., and Gould, K.L. (2015). The F-BAR Cdc15 promotes contractile ring formation through the direct recruitment of the formin Cdc12. J Cell Biol 208, 391–399.

Wu, J.Q., Kuhn, J.R., Kovar, D.R., and Pollard, T.D. (2003). Spatial and temporal pathway for assembly and constriction of the contractile ring in fission yeast cytokinesis. Dev Cell 5, 723–734.

Wu, J.Q., McCormick, C.D., and Pollard, T.D. (2008). Chapter 9: Counting proteins in living cells by quantitative fluorescence microscopy with internal standards. Methods Cell Biol 89, 253–273.

Wu, J.Q., and Pollard, T.D. (2005). Counting cytokinesis proteins globally and locally in fission yeast. Science 310, 310–314.

Wu, J.Q., Sirotkin, V., Kovar, D.R., Lord, M., Beltzner, C.C., Kuhn, J.R., and Pollard, T.D. (2006). Assembly of the cytokinetic contractile ring from a broad band of nodes in fission yeast. J Cell Biol 174, 391–402.

