## Supplemental Figures and Table for "Cooperation between Imp2p and Cdc15p imparts stiffness to the constricting contractile ring in fission yeast"

**SUPPLEMENTAL MATERIAL**

### Supplemental Figure 1

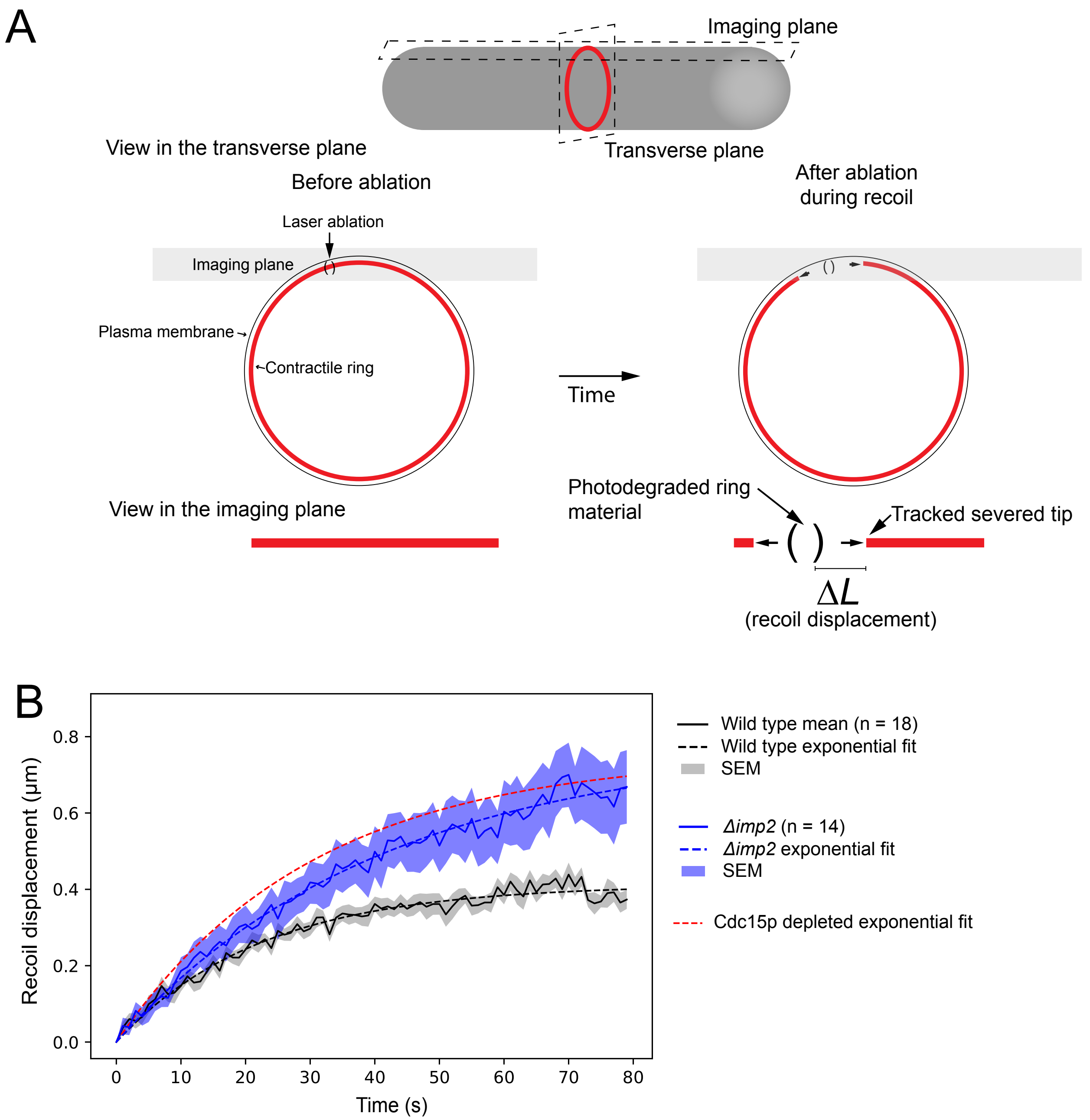


**Supplemental Figure 1. Imp2p and Cdc15p similarly impact the stiffness of the constricting contractile ring. A.** Diagram of the ablation of contractile rings. **B.** Graph of the displacement of severed tips in wild-type and $\Delta$*imp2* (as shown in Figure 1C) cells combined with the fit of the displacement from Cdc15p-depleted cells.

### Supplemental Figure 2

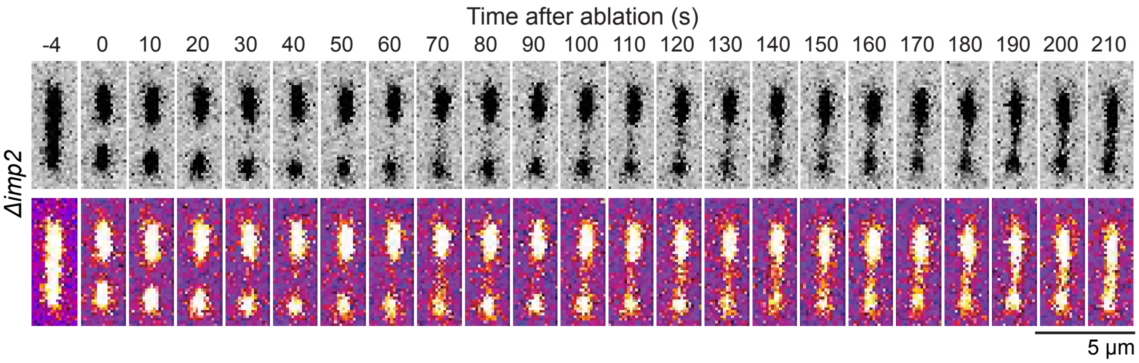


**Supplemental Figure 2. Severed contractile rings of Δ*imp2* cells heal by appearance of strands of mEGFP-Myp2p.** Example of severed Δ*imp2* cells where both severed tips are visible after ablation and after recoil. Material labeled with mEGFP-Myp2p gradually appears into focus across the gap ~ 70 s after ablation. Top row, inverted gray LUT. Bottom row, Fire LUT.

### Supplemental Figure 3

A

| | mEos3.2-Imp2p (27) | Imp2p-mEos3.2 (43) | Cdc15p-mEos3.2 (34) | mEos3.2-Cdc15p (51) |
|---|---|---|---|---|
| mEos3.2-Imp2p | - | | | |
| Imp2p-mEos3.2 | Y | - | | |
| Cdc15p-mEos3.2 | Y | Y | - | |
| mEos3.2-Cdc15p | Y | N | Y | - |

B

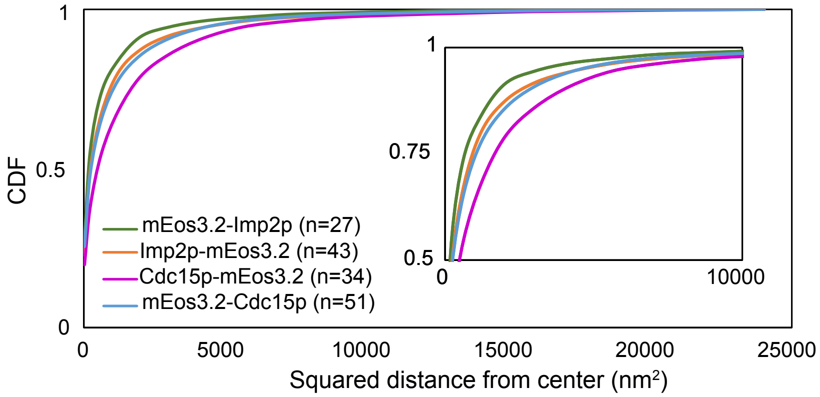


**Supplemental Figure 3. The molecular organization of Imp2p is significantly different from Cdc15p. A.** Table of KS tests used to determine whether the distribution of localizations was significantly different between pairs of proteins within a structure. Yes (Y), significant difference at $p < 0.005$. No (N), not significantly different. Sample size for each marker is in parentheses. **B.** Plots of the CDF for proteins measured in (A). Full CDF curve of data shown in Figure 3E. Inset, close-up of CDF curves.

### Table S1

| Strain | Genotype | Reference/Origin |
|---|---|---|
| **FPALM** | | |
| IRT201 | *h- kanMX6-Pcdc15-mEos3.2-cdc15 cdc25-22* | Laplante et al., 2016 |
| IRT206 | *h- cdc15-mEos3.2-kanMX6 cdc25-22 ade6-M210 ura4-D18* | Laplante et al., 2016 |
| CL812 | *h- imp2-mEos3.2-kanMX6 cdc25-22* | This work |
| CL893 | *kanMX6-Pimp2-mEos3.2-imp2 cdc25-22 ura4-D18* | This work |
| **Confocal** | | |
| #1723 | *bgs1Δ::ura4+ Pbgs1+-GFP-12A-bgs1-leu1+ leu1-32 ura4-D18 his3-D1* | Dr. Pilar Perez's lab. |
| CL174 | *h+ kanMX6-Pmyp2sh-mEGFP-myp2 ade6-M216 his3-D1 leu1-32 ura4-D18* | Laplante et al., 2015 |
| CL560 | *h- kanMX6-Pmyo2short-mEGFP-myo2 ade6-M210 his3-D1 leu1-32 ura4-D18* | Bellingham-Johnstun et al., 2021 |
| CL563 | *h+ kanMX6-Pmyo2short-mEGFP-myo2 sad1-RFP-kanMX6* | Bellingham-Johnstun et al., 2021 |
| CL586 | *h+ kanMX6-Pcdc15-mEGFP-cdc15* | Moshtohry et al., 2022 |
| CL607 | *h+ kanMX6-nmt81-mCherry-cdc15 kanMX6-Pmyo2short-mEGFP-myo2 sad1-RFP-kanMX6* | Moshtohry et al., 2022 |
| CL770 | *h+ imp2-mEGFP-kanMX6 ade6-M216 his3-D1 leu1-32 ura4-D18* | This work |
| CL814 | *Δimp2::natMX6 KanMX6-Pmyp2short-mEGFP-myp2 sad1-RFP-kanMX6* | This work |
| CL816 | *Δimp2::natMX6 KanMX6-Pmyo2short-mEGFP-myo2 sad1-RFP-kanMX6* | This work |
| CL912 | *cdc15-HALO-kanMX6 imp2-mEGFP-kanMX6* | This work |
| CL913 | *kanMX6-Pnmt81-mCherry-cdc15 imp2-mEGFP-kanMX6* | This work |
| CL914 | *Δimp2::natMX6 kanMX6-Pcdc15-mEGFP-cdc15* | This work |
| CL919 | *leu2::kanMX-Pcof1-mEGFP-Lifeact ura4-D18 leu1-32 ade6-M21X* | This work |
| CL931 | *Δimp2::natMX6 bgs1Δ::ura4+ Pbgs1+-GFP-12A-bgs1-leu1+ leu1-32 ura4-D18* | This work |
| CL933 | *Δimp2::natMX6 leu2::kanMX-Pcof1-mEGFP-Lifeact leu1-32 ura4-D18* | This work |
| **Calibration curve** | | |
| FY528 | *h+ ade6-M216 his3-D1 leu1-32 ura4-D18* | Forsburg lab. |
| SSP001 | *h+ mid1-mEGFP-kanMX6 leu1-32 ura4-D18 his3-D1 ade6-M210* | Saha et al., 2012 |
| JW1100 | *h+ spn1-mEGFP-kanMX6 ade6-M210 leu1-32 ura4-D18* | Wu et al., 2010 |
| KV344-1 | *h+ cdc12-3GFP-kanMX6 ade6-M216 leu1-32 ura4-D18 his3-D1* | Bellingham-Johnstun et al., 2021 |
| CL174 | *h+ kanMX6-Pmyp2sh-mEGFP-myp2 ade6-M216 his3-D1 leu1-32 ura4-D18* | Laplante et al., 2015 |
| CL537 | *h+ kanMX6-Prng2-mEGFP-rng2* | This work |
| CL560 | *h- kanMX6-Pmyo2short-mEGFP-myo2 ade6-M210 his3-D1 leu1-32 ura4-D18* | Bellingham-Johnstun et al., 2021 |
| CL586 | *h+ kanMX6-Pcdc15-mEGFP-cdc15* | Moshtohry et al., 2022 |

**Table S1. Table of strains used in this work.**
